# A modified dual preparatory method for improved isolation of nucleic acids from laser microdissected fresh-frozen human cancer tissue specimens

**DOI:** 10.1101/2024.08.02.606193

**Authors:** Danielle C. Kimble, Tracy J. Litzi, Gabrielle Snyder, Victoria Olowu, Sakiyah TaQee, Kelly A. Conrads, Jeremy Loffredo, Nicholas W. Bateman, Camille Alba, Elizabeth Rice, Craig D. Shriver, George L. Maxwell, Clifton Dalgard, Thomas P. Conrads

## Abstract

A central theme in cancer research is to increase our understanding of the cancer tissue microenvironment (TME), which is comprised of a complex and spatially heterogeneous ecosystem of malignant and non-malignant cells, both of which actively contribute to an intervening extracellular matrix. Laser microdissection (LMD) enables histology selective harvest of cellular subpopulations from the tissue microenvironment for their independent molecular investigation, such as by high-throughput DNA and RNA sequencing. Although enabling, LMD often requires a labor-intensive investment to harvest enough cells to achieve the necessary DNA and/or RNA input requirements for conventional next generation sequencing workflows. To increase efficiencies, we sought to use a commonplace dual preparatory (DP) procedure to isolate DNA and RNA from the same LMD harvested tissue samples. While the yield of DNA from the DP protocol was satisfactory, the RNA yield from the LMD harvested tissue samples was significantly poorer compared to a dedicated RNA preparation procedure. We identified that this low yield of RNA was due to incomplete partitioning of RNA in this widely used DP protocol. Here we describe a modified DP protocol that effectively partitions nucleic acids and results in significantly improved RNA yields from LMD harvested cells.

## Introduction

The use of laser microdissection (LMD) represents an important technology for collecting histology-resolved human tissue specimens for genomic, transcriptomic, and proteomic analyses. It is especially invaluable in cancer research, as tumor tissue can be readily separated from stroma or other cellular components, enabling studies of the tumor microenvironment ^1–4^. It evolved from early laser capture microdissection (LCM) technology that uses thermoplastic film coated caps and an infrared (IR) laser to adhere selected tissue area to the film ^1,3–6^. Subsequent technology allowed for the capture of selected tissue by ultraviolet (UV) laser ^2,7,8^ and collection by gravity, including the Leica LMD7 system used in this study ^5,9^. In this last iteration of LMD, thin tissue sections are mounted on membrane coated microscope slides, the tissue regions of interest (ROI) are cut with a UV laser and drop by gravity into a collection tube ^2,5^. This process allows for selective harvest of tissue segment ROI on the basis of histological morphology or as defined by immunohistochemical staining ^5^. A principal advantage of LMD over manually scraped tissue specimens is the ability to isolate specific tissue ROI or cell types of interest more precisely, aiding in spatially resolved tumor analyses, and with less contamination risk ^5,10^. Therefore, LMD is the ideal process to ensure higher sample purity; however, it takes significantly more time to harvest specimens by LMD as compared to macrodissection techniques such as manual scraping.

Another time and tissue consuming aspect of the sample processing workflow is nucleic acid isolation. Previous advances in methodology led to the development of both liquid extraction-based and spin column-based protocols for the simultaneous isolation of DNA and RNA from the same tissue sample ^11–13^. In particular, the Qiagen AllPrep DNA/RNA Kits allow for the simultaneous isolation of DNA and RNA from various tissue types using silica-based spin columns. This dual isolation from the same specimen collection saves precious tissue sample and reduces the microdissection burden compared to what is necessary for the isolation of DNA and RNA using single/dedicated preparatory procedures. Several studies have utilized the AllPrep DNA/RNA FFPE Kit to isolate nucleic acids from macrodissected formalin-fixed, paraffin-embedded (FFPE) tissue samples ^14–16^, while others have used the AllPrep DNA/RNA Micro Kit with macrodissected fresh frozen tissue or blood-based samples ^17,18^. Moreover, additional studies have reported the efficiency of the Qiagen AllPrep DNA/RNA Kit for IR-based LCM-procured fresh frozen tissue ^10^, and the use of the kit for RNA isolation from UV LMD-procured tissue for transcriptome profiling ^19^, but to our knowledge there have been no systematic evaluations of the efficiency of nucleic acid recovery from UV-based LMD-procured tissue using this widely used dual preparatory isolation method.

To decrease the overall processing time and labor investment incurred by LMD and nucleic acid isolation, we investigated the use of the Qiagen AllPrep DNA/RNA dual preparatory method for simultaneous DNA and RNA isolation from fresh frozen tumor tissue samples. Although the resulting DNA recoveries were sufficient in comparison to those obtained by a single preparatory method, the RNA recoveries were substantially lower than anticipated. Here we demonstrated the Qiagen AllPrep DNA/RNA isolation procedure results in the incorrect partitioning of RNA with the DNA isolate, an effect that selectively occurs with LMD procured fresh frozen tissue. To overcome this incorrect partitioning phenomenon, we developed and validated a modified isolation method enabling the selective extraction of RNA and DNA with high quality and quantity metrics with the addition of an AllPrep DNA “clean-up” column.

## Materials and Methods

### Tissue specimens and collection

Fresh frozen human cancer tissue specimens were obtained from women diagnosed with ovarian, peritoneal, and tubal cancers enrolled in the WCG IRB approved protocol #20110222 (Tissue and Data Acquisition Study of Gynecologic Disease) who underwent primary debulking surgery at Inova Fairfax Medical Campus. Written informed consent was obtained and the study was conducted in accordance with the Declaration of Helsinki.

Fresh frozen tumor specimens were embedded in optimal cutting temperature (OCT) compound and thin sectioned (4 µm) onto glass slides for generating diagnostic hematoxylin and eosin (H&E) sections for pathology review or sectioned (10 µm) onto polyethylene naphthalate (PEN) membrane slides (Leica Microsystems, Inc., Deerfield, IL) to harvest tissue by LMD for nucleic acid isolations. For membrane-coated slide testing, tissues were sectioned onto slides coated with polyethyleneimine (PEI), polyethylene terephthalate (PET), polyester (POL), or polyphenylene sulfide (PPS) (Leica Microsystems, Inc.). Slides were imaged and tumor-rich areas were annotated for collection with an Aperio AT2 slide scanner and Aperio ImageScope software (Leica Microsytems, Inc.). Approximately 65 mm^2^ of the selected sections was manually scraped using a scalpel or harvested by LMD (LMD7, Leica Microsystems, Inc.) into the appropriate buffer for each nucleic acid isolation kit and stored at −80 °C. To minimize degradation of RNA, tissue specimen slides remained at room temperature (RT) for LMD for no longer than 2 h.

### Original dual preparatory isolation of DNA and RNA

Dual preparatory (DP) isolation of DNA and RNA from scrape or LMD tissues was performed on tissue samples collected into 45 µL Buffer RLT Plus with 1% β-mercaptoethanol (BME, Sigma-Aldrich, St. Louis, MO, #M3148) using the AllPrep DNA/RNA Micro Kit (Qiagen Sciences, LLC, Germantown, MD, #80284) according to the manufacturer’s protocol in the “AllPrep® DNA/RNA Micro Handbook,” Simultaneous Purification of Genomic DNA and Total RNA from Microdissected Cryosections, including the optional DNase digestion. All reagents used are supplied with these kits unless otherwise stated. This will subsequently be referred to as the “original DP” protocol.

Briefly, the samples were thawed on ice and brought up to 350 µL with Buffer RLT Plus with BME. The tubes were vortexed for 30 s and briefly centrifuged. The sample was then added to an AllPrep DNA mini spin column and centrifuged for 30 s at 8000 x *g* (all centrifuge steps performed at RT). The column was stored at 4 °C, while the flow-through was used for RNA isolation. Next, 350 µL of 70% ethanol was added to the flow-through and mixed well by pipetting up and down 10 times. Then the sample was transferred to an RNeasy MinElute spin column, centrifuged for 15 s at 8000 x *g*, washed with 350 µL of Buffer RW1, and centrifuged for 15 s at 8000 x *g*. DNase digestion was performed by adding 80 µL of DNase solution (10 µL RNase-free DNase I stock (Qiagen #79254) plus 70 µL Buffer RDD (Qiagen #79254)) to the column and incubating for 15 min at RT. Then the column was washed with 350 µL Buffer RW1, centrifuged for 15 s at 8000 x *g*, washed with 500 µL Buffer RPE, and centrifuged for 30 s at 8000 x *g*. Next, 500 µL of 80% ethanol was added, and the column was centrifuged for 2 min at full speed (14000 x *g*), followed by a 5 min centrifugation step at full speed in a new collection tube to remove any remaining ethanol. Finally, the RNA was eluted into an Eppendorf DNA LoBind 1.5 mL microcentrifuge tube (Fisher Scientific, Waltham, MA, #13-698-791) by the addition of 17 µL RT RNase-free water to the column, incubated for 1 min at RT, then centrifuged for 1 min at full speed. Isolated RNA was then kept at −20 °C for short term storage or at −80 °C for long term storage.

Next, the AllPrep DNA mini spin column from above was removed from 4 °C, washed with 500 µL Buffer AW1, centrifuged for 30 s at 8000 x *g*, washed with 500 µL Buffer AW2, and centrifuged for 2 min at full speed. DNA was eluted by the addition of 40 µL of 70 °C Buffer EB to the column, incubated for 2 min at RT, and then centrifuged for 1 min at full speed. This was followed by two additional elution steps with 100 µL each of 70 °C RNase free water, incubated on the column for 2 min at RT, and centrifuged for 1 min at full speed. The final elution volume of 240 µL was reduced to 40 µL by vacuum centrifugation (CentriVap Concentrator, Labconco, Kansas City, MO). The isolated DNA was stored at −20 °C for short term storage or at −80°C for long term storage.

### Modified dual preparatory isolation of DNA and RNA

An additional AllPrep DNA column, acting as a “clean-up” column, was added to the beginning of the original DP protocol for all LMD tissue samples, which will hereafter be referred to as the “modified DP” protocol. Briefly, the samples containing LMD tissue were thawed on ice and bought up to a final volume of 350 µL with Buffer RLT Plus with BME. The tubes were vortexed for 30 s to homogenize the sample and briefly centrifuged. The sample was applied to the AllPrep DNA “clean-up” column, centrifuged for 30 s at 8000 x *g*, washed with 500 µL Buffer AW1, centrifuged for 30 s at 8000 x *g*, washed with 500 µL Buffer AW2, and centrifuged for 2 min at full speed. Subsequently, elution was performed by the addition of 40 µL of 70 °C Buffer EB to the column, followed by two elution steps of 100 µL each of 70 °C RNase free water, incubated for 2 min at RT and centrifuged for 1 min at full speed, resulting in a final elution volume of 240 µL. The volume was reduced to 40 µL by vacuum centrifugation. This elution contains the co-isolated DNA and RNA. Next, the sample was brought up to 350 µL with Buffer RLT Plus (no BME) and the full “original DP” isolation protocol was then performed as described above.

### Single preparatory DNA isolation

Tubes containing fresh frozen tissue specimens collected in Buffer ATL were removed from −80 °C, thawed on ice, and brought up to 360 µL with Buffer ATL. Lysis was performed by adding 40 µL of Proteinase K and incubating at 56 °C for 3 h with intermittent shaking. Single preparatory (SP) DNA isolation was performed using the QIAamp DNA Mini Kit (Qiagen, #51304) according to the manufacturer’s protocol, DNA Purification from Tissues found in the QIAamp DNA Mini and Blood Mini Handbook. DNA was eluted after a 10 min incubation at RT with 40 µL Buffer AE, followed by another 10 min incubation at RT with 160 µL nuclease-free water (Thermo Fisher Scientific, Inc., Waltham, MA, #AM9937) and reduced to 40 µL by vacuum centrifugation.

### Single preparatory RNA isolation

Tubes containing fresh frozen tissue specimens collected in Buffer RLT with 10% BME were removed from −80 °C, thawed on ice, and brought up to 300 µL with Buffer RLT with 10% BME. Single preparatory (SP) RNA isolation was performed using the RNeasy Micro Kit (Qiagen, #74004) per the manufacturer’s protocol, Purification of Total RNA from Microdissected Cryosections, including the on-column DNase digestion step, and eluted in 17 µL RNase-free water.

### Nucleic acid quantity and quality

Purity (260/280 ratio) was established spectrophotometrically (Nanodrop 2000 Spectrophotometer, Thermo Fisher Scientific, Inc.) and quantity was measured using the Qubit Broad Range (BR) or High Sensitivity (HS) kits, following the manufacturer’s instructions (Thermo Fisher, dsDNA HS (#Q32851), dsDNA BR (#Q32850), RNA HS (#Q32852), RNA BR (#Q10210)). Additional quality testing was completed using the 4200 Tapestation System (#G2991BA) and the genomic DNA ScreenTape (#5067-5365) and reagents (#5067-5366) to determine the DNA integrity numbers (DIN), or the RNA High Sensitivity ScreenTape (#5067-5579), Sample Buffer (#5067-5580), and Ladder (#5067-5581) to determine the RNA quality (RINe and DV200), following the manufacturer’s protocols (Agilent Technologies, Inc., Santa Clara, CA).

## Results

### RNA recovery using the original dual preparatory protocol

Motivated by the opportunity to gain efficiencies and spare the use of tissue, we sought to use a DP procedure to isolate tumor tissue DNA and RNA from the same LMD harvested tissue sample. Both LMD and manually procured (scalpel scraped) tissue specimens were processed using the original AllPrep DNA/RNA Micro Kit protocol (“original DP” protocol, Fig. 1) with a uniform input amount of 65 mm^2^ tissue per sample. Unexpectedly, we found that the recovery of RNA in the RNA isolate tube from LMD procured tissue was diminishingly low with only an average of 124 and 131 ng recovered for specimens A047 and A067, respectively, compared to their scraped counterparts with approximately 10-fold higher values (Fig. 2A and B). Using fluorometric assays uniquely selective for either DNA or RNA, we identified that RNA was co-isolating with DNA in the DNA isolate tube for the LMD procured samples, with an average of 88-90% of the total RNA retained in the DNA isolate (Fig. 2A and B). When processed with the same DP method, the scraped samples resulted in the correct partitioning of RNA and DNA, with an average of 82-86% of the total RNA yield recovered in the RNA isolate tube (Fig. 2A and B). The DNA recovery was not adversely affected by the tissue procurement method (data not shown). This result led us to evaluate various steps in the LMD and nucleic acid isolation workflows to determine the cause of the retention of RNA in the DNA isolate.

**Figure 1.**
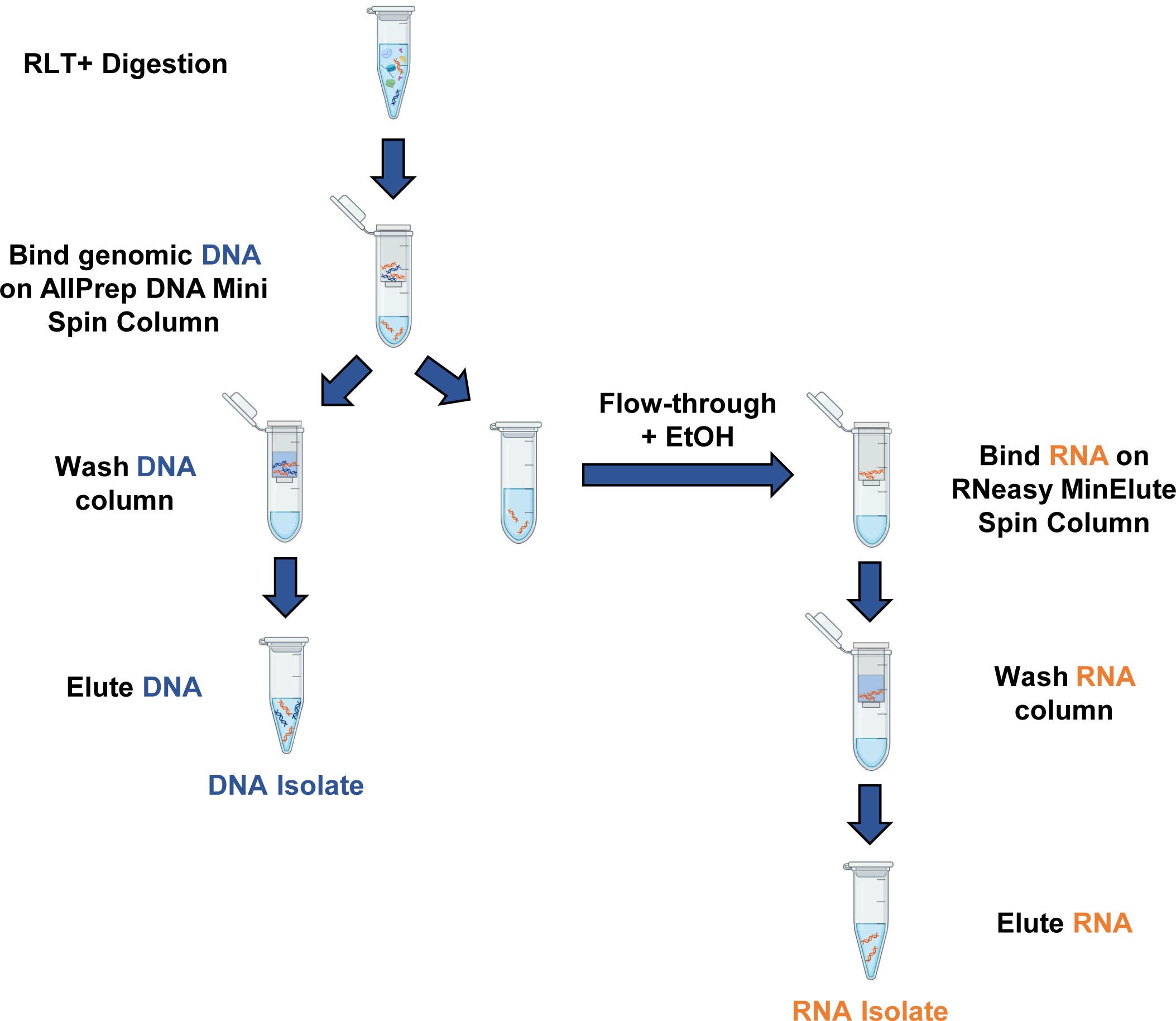
Workflow of the original dual preparatory protocol. The original AllPrep DNA/RNA Micro Kit based on the manufacturer’s protocol. After sample digestion in Buffer RLT Plus, DNA is bound to an AllPrep DNA column while the flow-through is supplemented with 70% EtOH and processed on an RNeasy column to isolate the RNA. DNA is then washed and eluted off the DNA column. Blue and orange strands represent DNA and RNA, respectively. The figure was created with BioRender.com.

**Figure 2.**
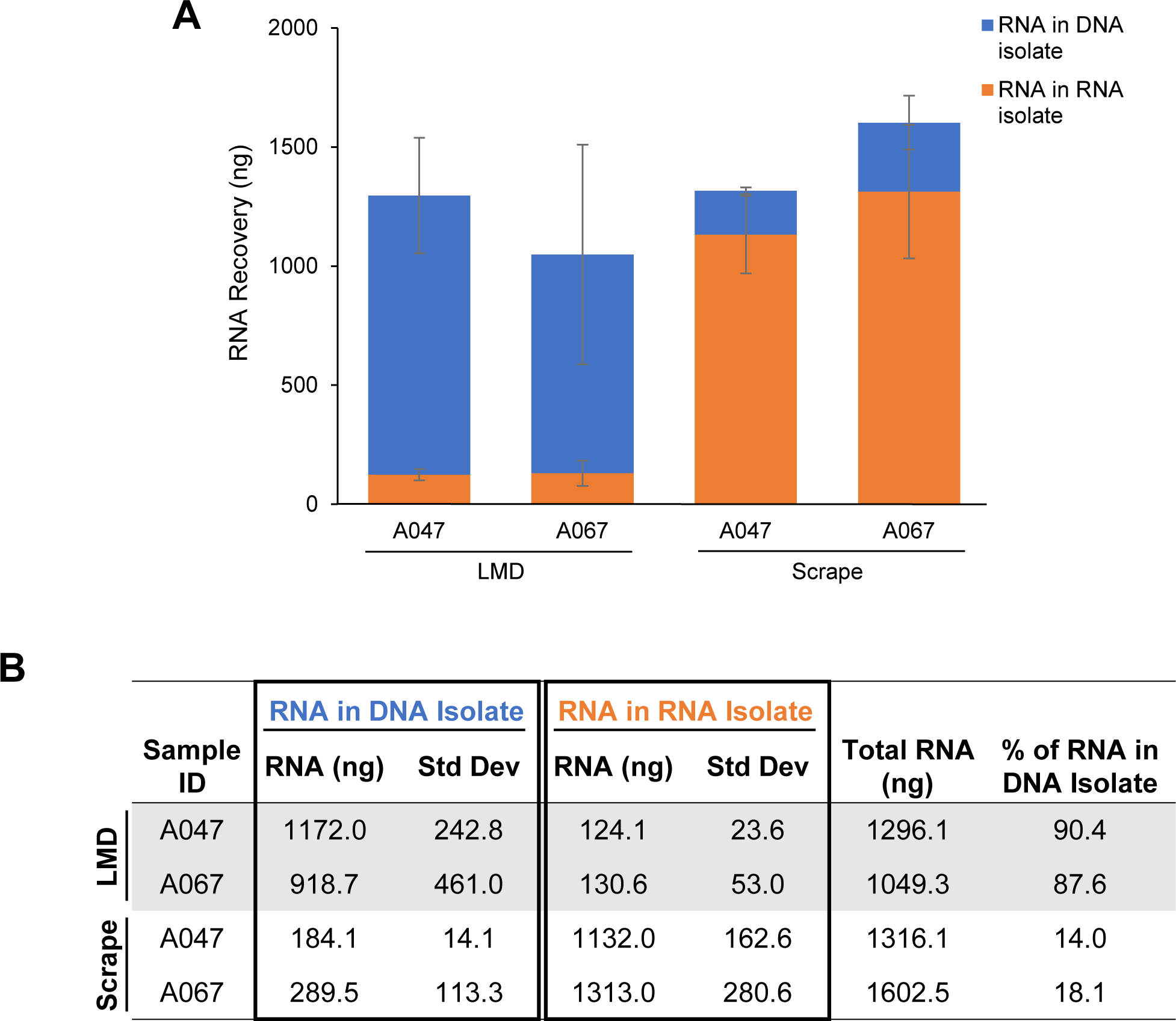
Initial evaluation of RNA recovery in LMD vs. scraped tissues using the original DP method. **(A and B)** Average total amount of RNA recovered using ∼65 mm^2^ of either LMD or scrape-procured tissue samples processed by the original DP protocol. RNA recovery was measured by Qubit in both the RNA isolate tubes (orange) and DNA isolate tubes (blue). Data for each unique sample ID is represented as the average total ng of RNA isolated from biological replicates (*n* = 3-4). Standard deviation bars are shown. Specificity of the Qubit RNA assay was confirmed by control DNA and RNA (data not shown).

### Troubleshooting the incorrect partitioning of RNA

#### Slide handling and membrane characteristics

Fresh frozen tissue mounted on polyethylene naphthalate (PEN) membrane coated slides were washed with 100% ethanol to remove condensate immediately prior to LMD. We hypothesized that residual ethanol may interfere with the RNA column binding properties. To test this, we collected LMD tissue with and without the ethanol wash step, as well as scraped tissue as the control. Eliminating the ethanol wash step had no effect on the RNA isolation, as greater than 80% of RNA remained co-isolated with the LMD procured DNA samples regardless of EtOH status (Supplementary Fig. S1).

We next looked at the effect of various slide compositions to determine if the co-isolation of RNA and DNA was due to an aspect specifically related to polyethylene naphthalate (PEN). Tissue sections were mounted on slides coated with either PEN, polyethyleneimine (PEI), polyethylene terephthalate (PET), polyester (POL), or polyphenylene sulfide (PPS), a total area of ∼65 mm^2^ was harvested by LMD with biological replicates, and the original DP protocol was followed to isolate DNA and RNA. Samples from all membrane types showed similar results, with a large proportion of RNA (78-91%) co-isolating with the DNA (Fig. 3A and B).

**Figure 3.**
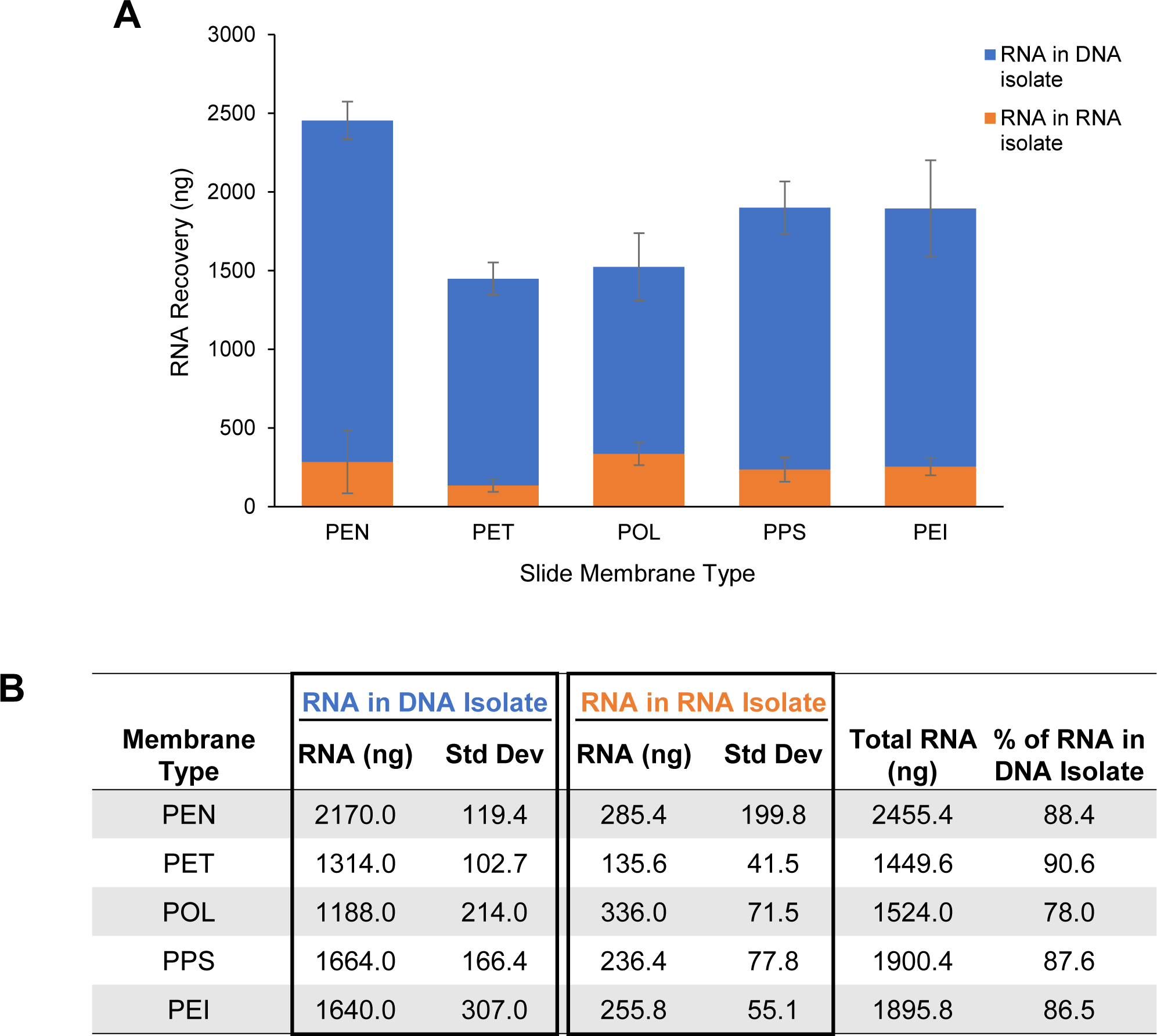
RNA recovery by microscope slide membrane type. **(A and B)** Recovery from ∼65 mm^2^ of specimen A067 LMD-procured tissue sections mounted on microscope slides coated with either polyethylene naphthalate (PEN), polyethylene terephthalate (PET), polyester (POL), polyphenylene sulfide (PPS), or polyethyleneimine (PEI). RNA was isolated with the original DP protocol, and recovery was measured by Qubit in both the RNA isolate tubes (orange) and DNA isolate tubes (blue). Bars represent the average total ng of RNA recovered from biological replicates (*n* = 4). Standard deviation bars are shown.

#### Comparison of SP and DP protocol RNA recovery

Previously, nucleic acids from fresh frozen tissue specimens procured via three methods, either LMD, scraping, or scrolling, were isolated with the QIAamp DNA Mini Kit (DNA SP protocol). Tissue scrolls were harvested directly off OCT blocks into a tube, with no PEN membrane present. All fresh frozen tissue specimens were collected from similar tissue types, and nucleic acids were isolated using the original DP protocol with the AllPrep DNA/RNA Micro Kit. We revisited these nucleic acid isolations and quantified the RNA in the DNA isolate tubes for both the SP and DP methods. Due to varied tissue input amounts, the ratio of RNA:DNA in the DNA isolate was used to enable comparisons. The original DP isolations from scraped samples had the lowest level of RNA in the DNA isolate tubes, with an average RNA:DNA ratio in the DNA isolate tube of 0.12, confirming our initial findings of effective nucleic acid partitioning for scraped samples (Fig. 4A and B). The RNA yield in the SP DNA isolate from tissue scrolls was also low, with an average RNA:DNA ratio of 0.28. For both the SP and DP LMD-procured samples, high RNA:DNA ratios in the DNA isolate averaged 0.89 and 1.20 respectively, indicating improper partitioning and co-isolation of the RNA with the DNA (Fig. 4A and B). Surprisingly, the scrape procured samples prepared with a DNA SP procedure resulted in a high RNA:DNA ratio in the DNA isolate as well, with an average of 0.81 (Fig. 4A and B). One striking difference between the SP and DP protocols is that cell lysis includes a 3 hr incubation at 56 °C for the SP method (heat), whereas only vortexing at room temperature is used in the DP method (no heat) (Fig. 4B). Because we observed effective RNA partitioning with the DP protocol (no heat) from scraped PEN membrane slide mounted tissue and with the SP method (heat) from tissue scrolls (no PEN), we speculated that heating, whether from incubation in the lysis step of the SP method or from UV laser-induced heating during the procurement of LMD tissue, in combination with the membrane-coated slides, is adversely impacting RNA partitioning on the silica-based spin columns used in these isolation procedures.

**Figure 4.**
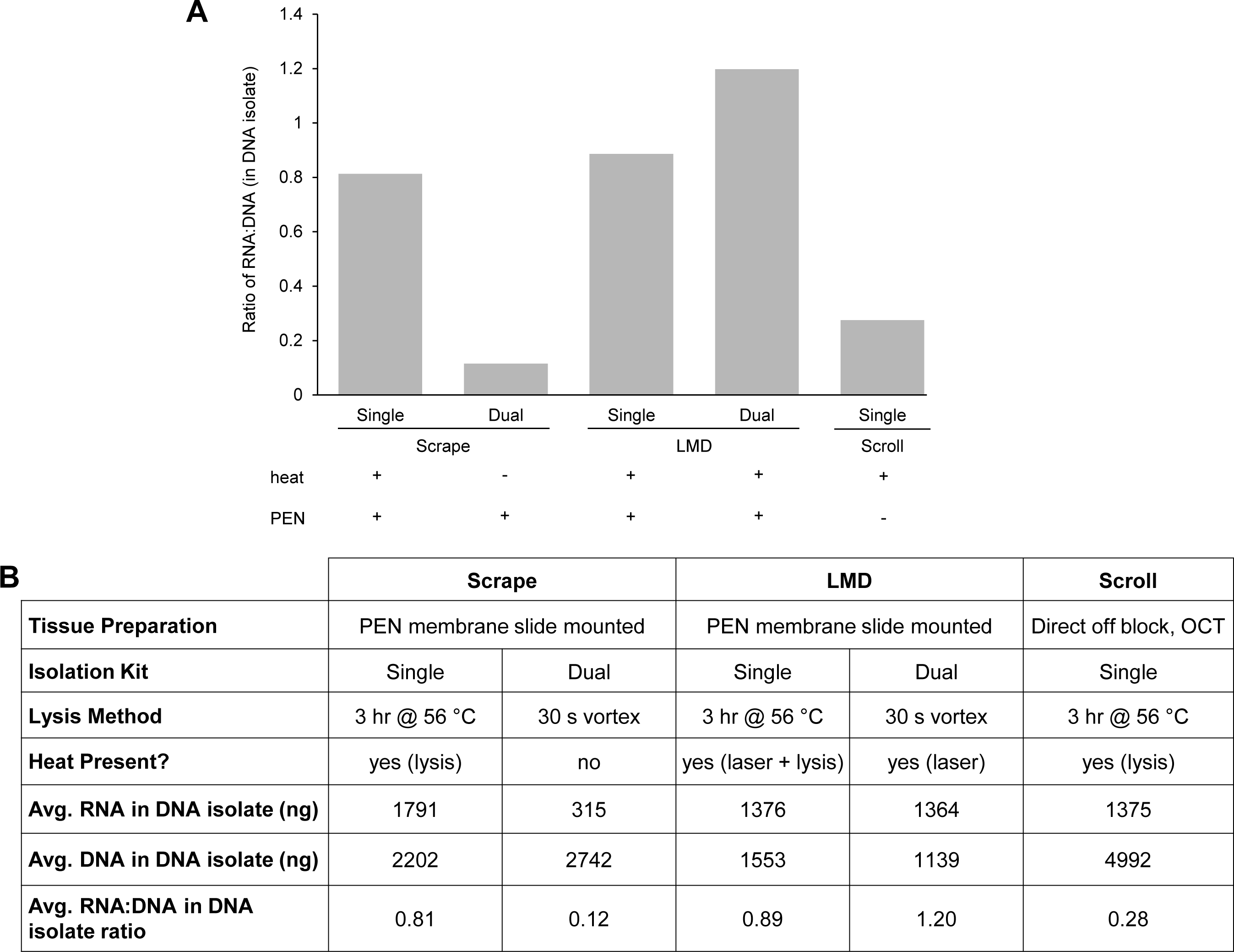
Comparison of SP and original DP protocol RNA recovery in DNA isolate. Fresh frozen tissue mounted on PEN membrane slides was harvested by either manually scraping or by LMD. Alternatively, tissue scrolled directly off an OCT block was used as a control. DNA was initially isolated with the QIAamp DNA Mini Kit protocol (single prep, SP) and later repeated using the original AllPrep DNA/RNA Mini Kit protocol (dual prep, DP). **(A)** DNA and RNA were quantified by Qubit and the result is displayed as a ratio of the RNA to the DNA, in the DNA isolate tube. Presence (+) or absence (-) of a heat source or PEN membrane for each method is indicated. **(B)** Comparison of nucleic acid metrics for the various tissue collection and lysis methods. For SP scrapes, DP scrapes, and SP LMD, *n* = 10 (each a unique sample ID); for SP scroll, *n* = 9 (each a unique sample ID); for DP LMD, *n* = 12 (6 biological replicates for 2 unique sample IDs).

#### Effect of heat from LMD UV laser and PEN membrane

To test potential effects from the heat of the LMD microscope UV laser in conjunction with PEN membrane-coated slides on RNA partitioning, we laser microdissected the equivalent of 65 mm^2^ “blank” PEN membrane elements and added these directly into Buffer RLT Plus with BME along with the immediate addition of 65 mm^2^ of scraped fresh frozen tissue. Using the original DP isolation method, the results of this experiment recapitulated those from direct LMD procured tissue, where we observed extremely low RNA yields in the RNA isolate tube and high RNA recovery in the DNA isolate tube (Fig. 5A and B). Conversely, freezing the blank PEN elements at −20 °C in Buffer RLT Plus with BME first prior to adding to the scraped tissue resulted in increased RNA partitioning in the RNA isolate (Fig. 5A and B). Only 32.3% of the total RNA remained in the DNA isolate when frozen elements were added (Scrape + PEN_F), as compared to 78.8% when the elements were added immediately without freezing (Scrape + PEN) (Fig. 5A and B). These results suggest that the immediate interaction between laser-mediated heated PEN membrane and tissue negatively impacts partitioning of RNA and results in co-isolation of RNA with DNA in the DP procedure.

**Figure 5.**
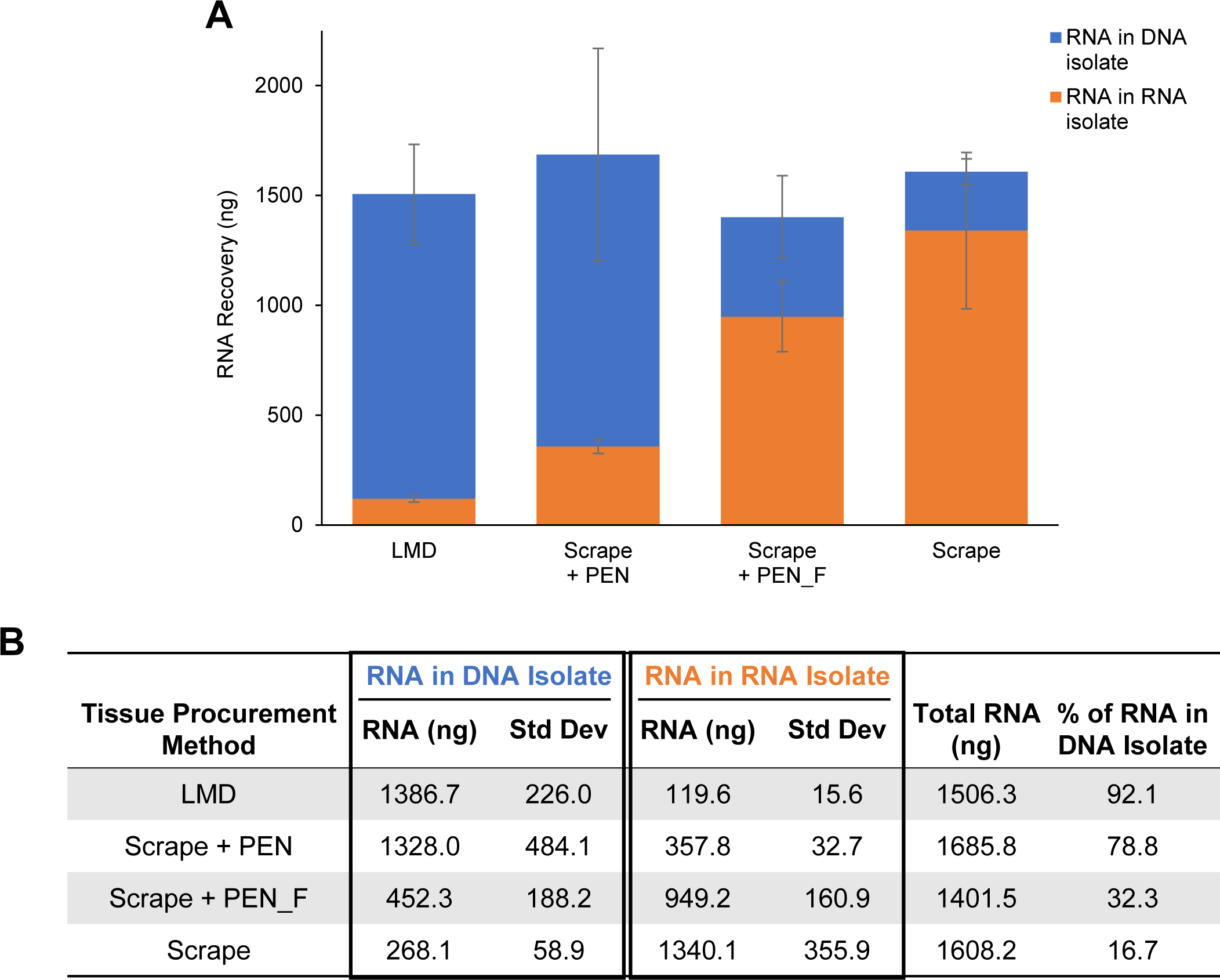
Effect of laser heat and membrane coating on RNA recovery. **(A and B)** Approximately 65 mm^2^ of LMD-procured blank PEN membrane elements were added to scrape-procured tissue from specimen A067 either immediately after LMD (Scrape + PEN) or after being frozen at −20°C (Scrape + PEN_F). LMD-procured and scrape-procured tissue on PEN membrane coated slides were used as controls. RNA was isolated with the original DP method and quantified by Qubit in the RNA isolate tubes (orange) and DNA isolate tubes (blue). Data is displayed as an average of biological replicates (*n* ≥ 3). Standard deviation bars are shown.

### Adaptation and assessment of modified DP protocol

After ruling out multiple variables related to histology preparation and LMD microscope slide membrane composition, we identified that laser-mediated heating of the membrane on the microscope slide during the LMD procurement process appeared to be the cause of improper partitioning of RNA. To resolve the incorrect partitioning, we tested the addition of an extra AllPrep DNA column to act as a “clean-up” step (“modified DP” protocol, Fig. 6). The digested tissue lysate was passed through the AllPrep DNA “clean-up” column, and then the eluate was processed using the original DP protocol with a fresh AllPrep DNA mini spin column. LMD procured tissue samples on PEN membrane slides processed by this modified DP protocol resulted in most of the RNA partitioned into the RNA isolate tube, and ≤ 25% being co-isolated with the DNA (Fig. 7). Using the modified DP protocol for LMD-procured tissue resulted in a 3 to 4-fold increase in the RNA recovered in the RNA isolate tube compared to using the original DP protocol (Fig. 7A and B). The amount of isolated RNA using this modified DP protocol was comparable to the amount of RNA isolated from the corresponding samples using the SP RNeasy Mini Kit (Supplementary Fig. S2, B). RNA quality as measured by RINe and DV200 values was not adversely affected by the protocol modification, as an average RINe value of 7.6 and a DV200 of 98.2% were obtained, compared to 6.9 and 93.1% for the SP RNA protocol (Supplementary Fig. S2, C). Subsequent RNA sequencing of RNA isolated from both protocols determined that there were no effects of the modified DP protocol on sequence data (data not shown).

**Figure 6.**
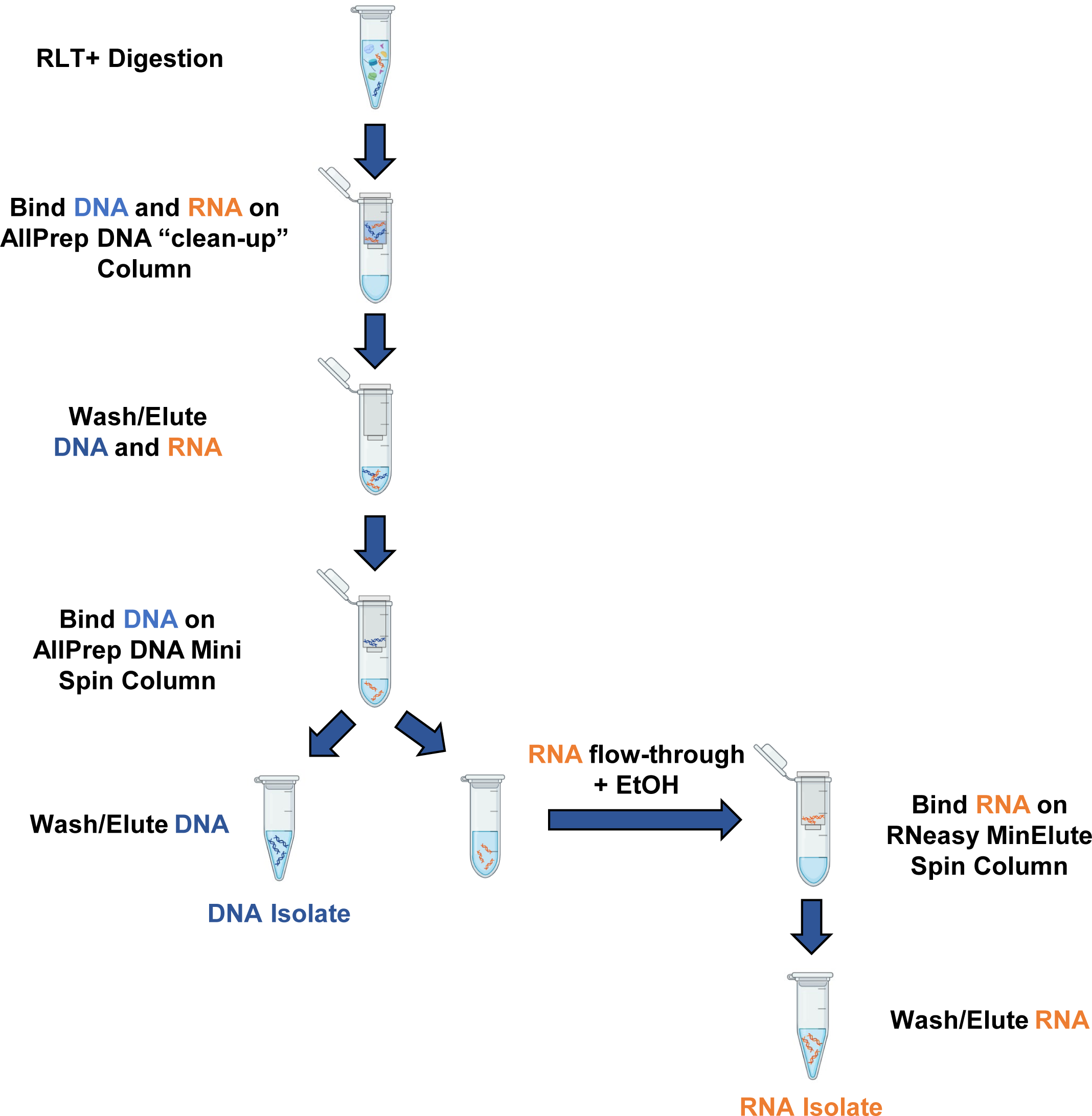
Workflow of the modified dual preparatory protocol. The modified version of the protocol, specific for LMD-procured samples from membrane-coated slides, which includes an extra AllPrep DNA column as a “clean-up” step after sample digestion. The eluate from the AllPrep DNA “clean-up” column contains both DNA (blue) and RNA (orange), which are then separated by the second AllPrep DNA column following the original protocol. The figure was created with BioRender.com.

**Figure 7.**
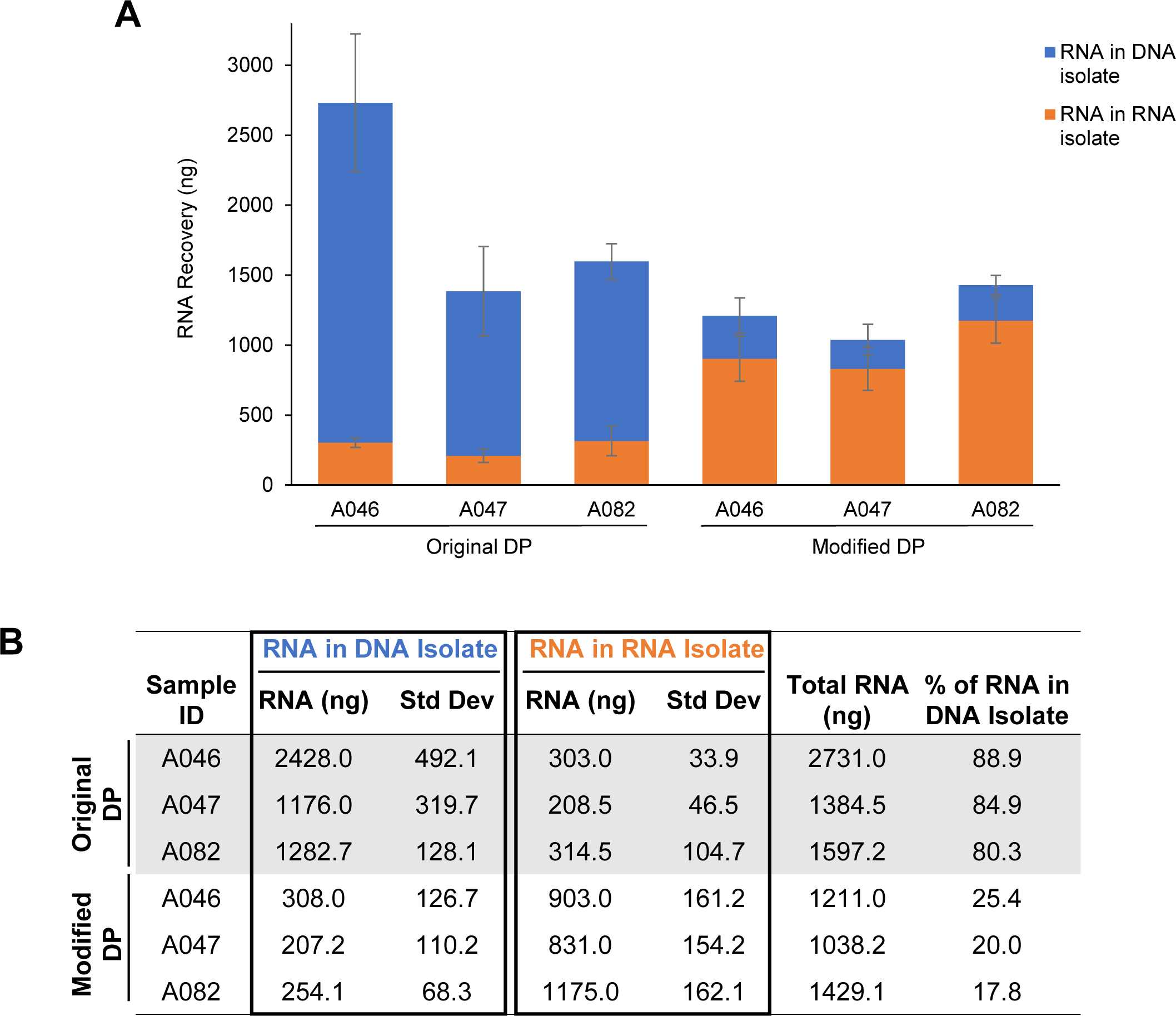
Improved RNA partitioning in LMD-procured tissue with the modified DP protocol. **(A and B)** The average total RNA recovered with ∼65 mm^2^ of LMD-procured tissue from specimens A046, A047, and A082 using either the original or modified DP protocol. RNA recovery was quantified by Qubit in both the RNA isolate tubes (orange) and DNA isolate tubes (blue). Data is displayed as an average of biological replicates (*n* = 3), and standard deviation bars are shown.

## Discussion

Due to the significant heterogeneity in the tumor microenvironment, laser microdissection has become a useful tool by enabling the histology selective harvest of cellular populations for molecular investigations. Conventional nucleic acid sequencing often requires a labor-intensive investment to harvest sufficient amounts of cells by LMD. Utilizing one tissue collection for both DNA and RNA reduces the chance of specimen handling errors and conserves time, as it would take twice as long to harvest tissue for single isolations of DNA and RNA. Dual isolation of DNA and RNA from a singular collected tissue sample is beneficial as tumor specimens can be limited in availability, and this allows for the remaining valuable tissue to be utilized for other molecular analyses (i.e., proteomics). Additionally, results from DNA and RNA sequencing will have higher correlations as the nucleic acids were isolated from the exact same tissue and cellular composition.

We tested the AllPrep DNA/RNA Micro Kit on in-house tumor samples following the manufacturer’s protocol and discovered the improper partitioning of RNA isolated from LMD-procured tissue. After eliminating variables associated with slide handling and membrane composition, we found the application of heat to membrane-coated slides to be the issue. Interestingly, the heat applied in the lysis step of the SP DNA isolation protocol for scrape-procured tissue on PEN membrane slides acted in the same manner as heat introduced by the LMD UV laser in the DP protocol for LMD-procured tissue on PEN membrane slides, resulting in RNA in the DNA isolate tubes (Fig. 4). We have provided evidence that this appears to be due to an immediate interaction between the tissue and laser-mediated heating of the polymer membrane on the microscope slide (Fig. 5). This occurred regardless of membrane composition type (Fig. 3).

We speculate that laser-mediated heating of LMD microscope slide membranes liberates an unknown chemical moiety that prevents partitioning of RNA from DNA in the DP procedure as designed by the manufacturer. To our knowledge, there have been no previous reports of this phenomenon of laser-mediated heating of LMD membrane-coated microscope slides interfering with RNA isolation. A possible explanation could be that a chemical moiety or altered characteristic such as chemical charge is causing either the RNA to bind to DNA or directly to the column. Further elucidating the mechanism of the co-isolation is limited by the fact that the composition of AllPrep DNA columns is proprietary, other than that they are silica based. Additional research would be necessary to determine the exact chemical change occurring.

To achieve correct RNA partitioning, we designed and validated a modified protocol with an additional “clean-up” column for LMD-procured tissue to allow for the separation of the RNA from the DNA, without any reductions in nucleic acid quality. We hypothesize that the effects of the chemical moiety are neutralized by this “clean-up” column, thereby allowing correct nucleic acid partitioning when the eluate is processed with the second AllPrep DNA column. Our efforts to re-design the manufacturer’s protocol for DP preparation of nucleic acids from LMD procured tissue has allowed for the continued use of DP isolation toward increased efficiencies of investment of time and tissue.

## Author contributions

Danielle C. Kimble (Investigation [supporting], Writing-original draft [lead], Writing-review and editing [equal]), Tracy J. Litzi (Conceptualization [equal], Investigation [equal], Methodology [lead], Writing-review and editing [equal]), Gabrielle Snyder (Investigation [equal], Methodology [equal], Writing-review and editing [equal]), Kelly A. Conrads (Investigation [supporting], Writing-review and editing [equal]), Jeremy Loffredo (Investigation [supporting], Methodology [lead], Writing-review and editing [supporting]), Nicholas W. Bateman (Investigation [supporting], Writing-review and editing [equal]), Camille Alba (Investigation [supporting], Methodology [supporting], Writing-review and editing [supporting]), Elizabeth Rice (Investigation [supporting], Methodology [supporting], Writing-review and editing [supporting]), Craig D. Shriver (Funding acquisition [lead], Writing—review and editing [supporting]), Clifton Dalgard (Investigation [supporting], Writing—review and editing [supporting]), Thomas P. Conrads (Conceptualization [lead], Formal analysis [lead], Funding acquisition [lead], Project administration [lead], Supervision [lead], Writing—review and editing [equal]).

## Disclaimer

The views expressed in this article are those of the author(s) and do not necessarily reflect the official policy or position of the Uniformed Services University of the Health Sciences (USUHS), Department of the Navy, Department of the Air Force, Department of the Army, Department of Defense, or the United States Government. Mention of trade names, commercial products, or organizations does not imply endorsement by the U.S. Government. The author(s) are military service members. This work was prepared as part of their official duties. Title 17 U.S.C. 105 provides that ‘Copyright protection under this title is not available for any work of the United States Government.’ Title 17 U.S.C. 101 defines a United States Government work as a work prepared by a military service member or employee of the United States Government as part of that person’s official duties.

## Funding

This work was funded in part by awards HU0001-16-2-0006, HU0001-19-2-0031, HU0001-20-2-0033, and HU0001-21-2-0027, and HU0001-22-2-0016 from the Uniformed Services University of the Health Sciences from the Defense Health Program to the Henry M Jackson Foundation for the Advancement of Military Medicine Inc. in support of the Gynecologic Cancer Center of Excellence Program.

## Conflict of interest statement

T.P.C. is a ThermoFisher Scientific, Inc SAB member and receives research funding from AbbVie.

---

**Supplementary Figure S1.**
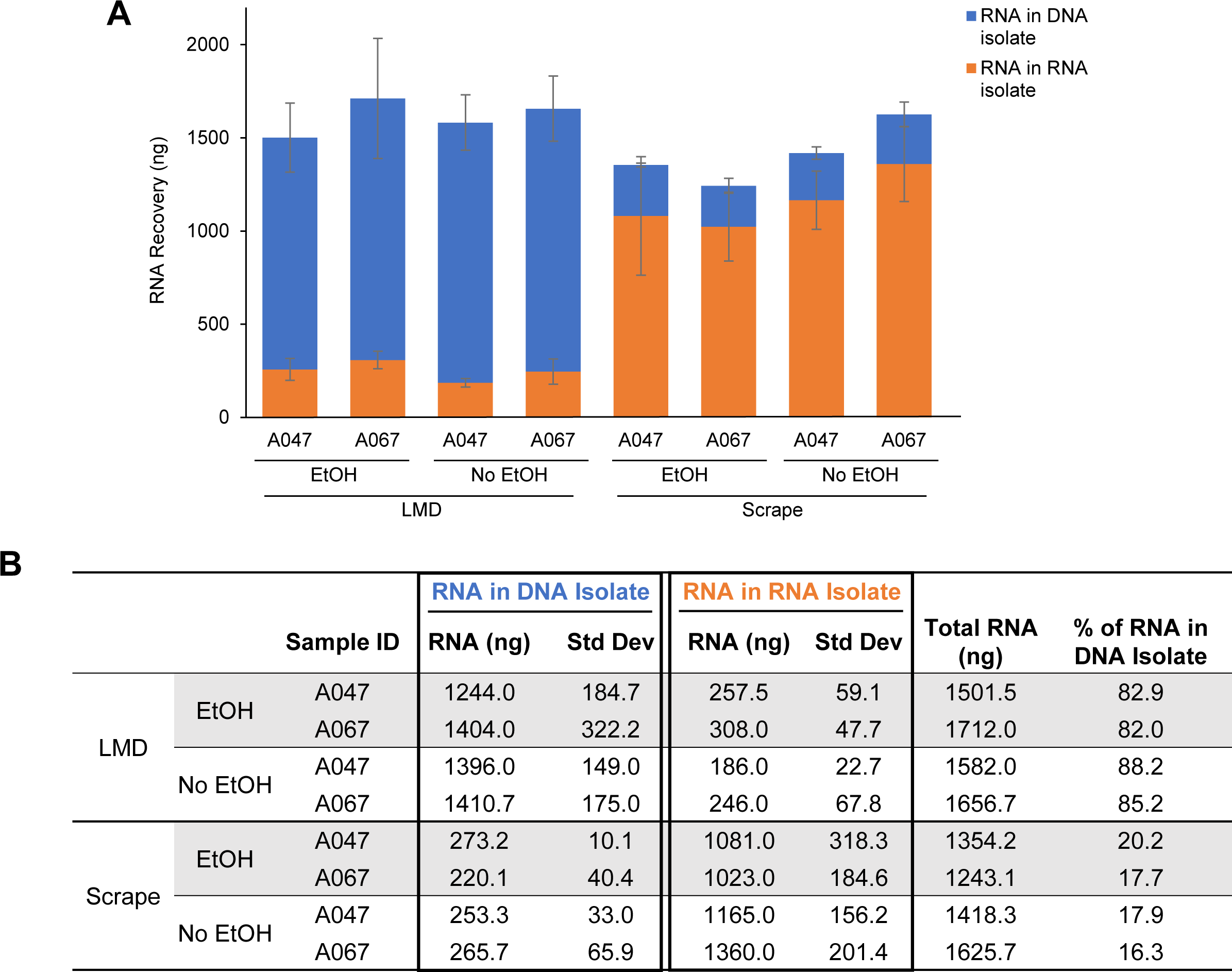
Effect of ethanol on RNA yield in LMD and scraped tissues. **(A and B)** Fresh frozen tissue specimens mounted onto PEN membrane-coated microscope slides were washed with 100% EtOH to remove condensate prior to harvesting by LMD. Slides were either used after this EtOH wash step (EtOH; standard protocol) or without an EtOH wash step and brought to room temperature first to remove residual condensation (No EtOH) prior to procurement by LMD or manually scraping. RNA was isolated with the original DP protocol and quantified by Qubit in both the RNA isolate tubes (orange) or DNA isolate tubes (blue). Bars represent the average total ng of RNA recovered from biological replicates (*n* = 3). Standard deviation bars are shown.

**Supplementary Figure S2.**
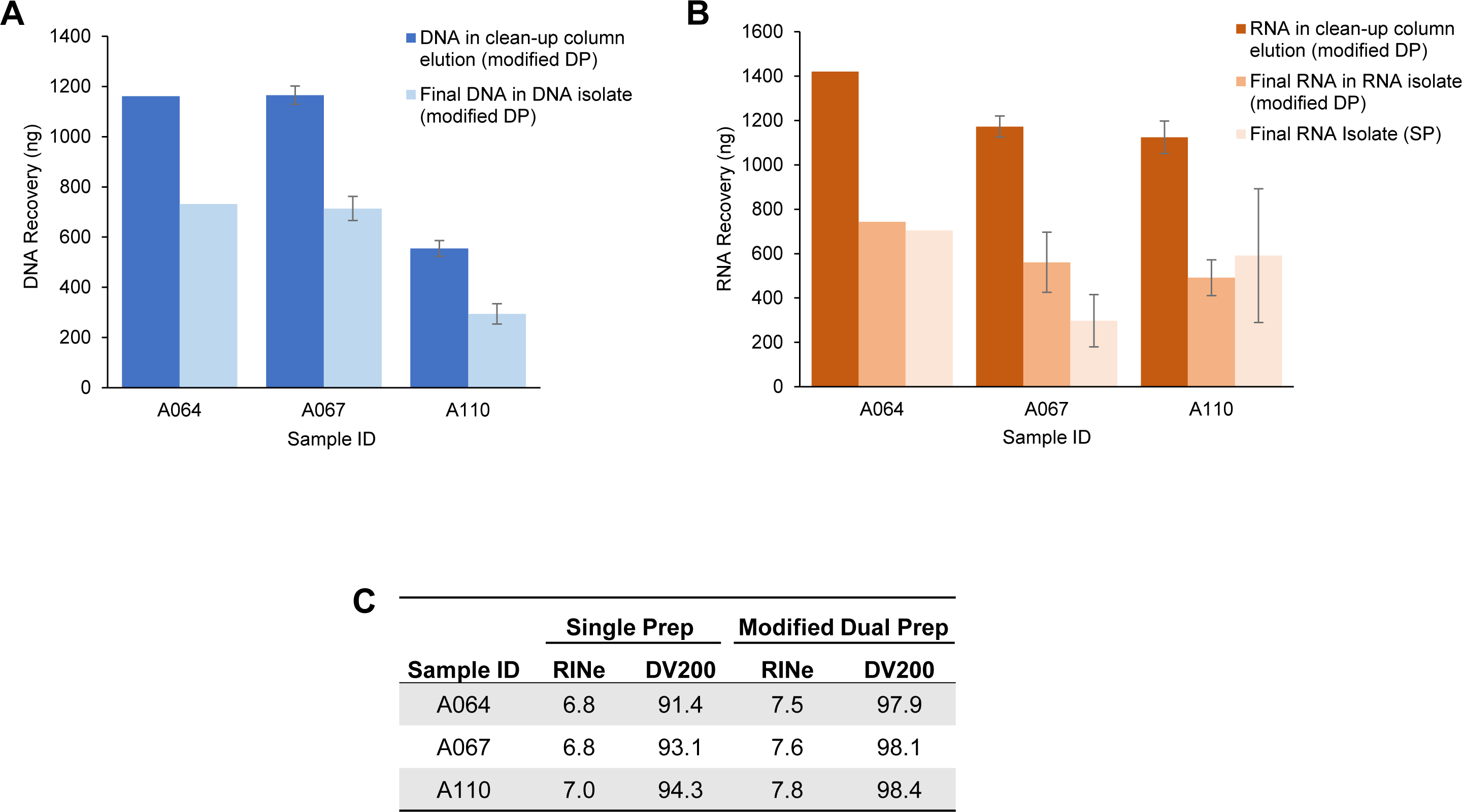
Efficiency and quality metrics of nucleic acid isolations. **(A)** Efficiency of DNA recovery with the modified DP protocol. The average total DNA recovered in the flow-through of the AllPrep DNA “clean-up” column compared to the average total DNA in the final DNA isolate as measured by Qubit (A064, *n* = 1; A067 and A110, *n* = 2). Standard deviation bars are shown. **(B)** Efficiency of RNA recovery with the RNA SP and modified DP protocols. The average RNA recovery in the flow-through of the AllPrep DNA “clean-up” column (RNA in clean-up column elution, modified DP) compared to the final RNA isolate (final RNA in RNA isolate, modified DP), as measured by Qubit (A064, *n* = 1; A067 and A110, *n* = 2). The average RNA recovery in the final RNA isolate of the SP protocol is also included. Standard deviation bars are shown. **(C)** Average RINe values and average percent of RNA fragments greater than 200 nucleotides (DV200) of the RNA recovered in the final RNA isolate tubes with the SP and modified DP methods (*n* ≥ 5). Data is an average of biological replicates from A064, A067, and A110.

## Notes

### Summary of Updates

Nicholas Bateman's name was misspelled in the initial version of this paper's submission.

